# Invasion fitness, inclusive fitness, and reproductive numbers in heterogeneous populations

**DOI:** 10.1101/036731

**Authors:** Laurent Lehmann, Charles Mullon, Erol Akçay, Jeremy Van Cleve

**Affiliations:** Department of Ecology and Evolution, University of Lausanne, Switzerland.; Department of Biology, University of Pennsylvania, USA.; Department of Biology, University of Kentucky, USA.

**Keywords:** growth rate, invasion fitness, inclusive fitness, reproductive number, invadability

## Abstract

How should fitness be measured to determine which phenotype or “strategy” is uninvadable when evolution occurs in subdivided populations subject to local demographic and environmental heterogeneity? Several invasion fitness measures, such as basic reproductive number, lifetime dispersal success of a local lineage, or inclusive fitness have been proposed to address this question, but the relationships between them and their generality remains unclear. Here, we ascertain uninvadability (all mutant strategies always go extinct) in terms of the growth rate of a mutant allele arising as a single copy in a population. We show from this growth rate that uninvadability is equivalently characterized by at least three conceptually distinct invasion fitness measures: (i) lineage fitness, giving the average personal fitness of a randomly sampled mutant lineage member; (ii) inclusive fitness, giving a reproductive value weighted average of the direct fitness cost and relatedness weighted indirect fitness benefits accruing to a randomly sampled mutant lineage member; and (iii) three types of reproductive numbers, giving lifetime success of a local lineage. Our analysis connects approaches that have been deemed different, generalizes the exact version of inclusive fitness to class-structured populations, and provides a biological interpretation of selection on a mutant allele under arbitrary strength of selection.

## Introduction

It is well established (if perhaps unwelcome) that in general adaptiveness is not increased by short-term evolution (Moran, 1964; Eshel, 1991; Ewens, 2004). In contrast, when long-term evolution can be described by a substitution process where a population transitions from one fixed allele to another through the recurrent invasion of mutant alleles, the population may eventually evolve to an uninvadable state (i.e., a state that is resistant to invasion by any alternative strategy, Eshel, 1991, 1996; Hammerstein, 1996; Weissing, 1996; Van Cleve, 2015). An uninvadable strategy is “optimal” among a specified set of alternatives because it maximizes the growth rate of the underlying coding gene when the gene is rare (Eshel, 1991,1996; Hammerstein, 1996; Weissing, 1996). Uninvadable strategies are thus adaptations *(sensu* Reeve and Sherman, 1993) and evolutionary invasion analysis has become a very successful approach to understand theoretically long term phenotypic evolution (e.g., Maynard Smith, 1982; Eshel and Feldman, 1984; Parker and Maynard Smith, 1990; Charlesworth, 1994; Metz et al., 1996; Ferrière and Gatto, 1995; McNamara et al., 2001; Lion and van Baalen, 2007; Metz, 2011; van Baalen, 2013).

When a mutant allele arises as a single copy in a population, its growth rate, *ρ*, determines in general whether the mutant allele will eventually go extinct or survive (Tuljapurkar, 1989; Metz et al., 1992; Rand et al., 1994; Charlesworth, 1994; Ferrière and Gatto, 1995; Caswell, 2000). Intuitively, the growth rate is a gene-centered measure of evolutionary success *(sensu* Dawkins, 1978). Technically, the growth rate is the dominant eigenvalue of a matrix determining the transitions between the different states in which the mutant allele can reside and describes the growth of a typical trajectory of the mutant lineage since its appearance as a single copy(Tuljapurkar, 1989; Tuljapurkar et al., 2003; Caswell, 2000; Ferrière and Gatto, 1995). Since evolutionary biologists often try to understand adaptations in terms of the fitness properties exhibited by individuals, such as survival and fecundity, it is important to understand the exact interpretation of the growth rate in terms of individual-centered fitness components. Interpreting the growth rate this way seems clear in panmictic populations. In the absence of genetic conflict within individuals, maximizing the growth rate amounts to maximizing the personal (lifetime) fitness of an individual, which is determined by its survival and fecundity schedules in stage-structured populations (Eshel and Feldman, 1984; Hammerstein, 1996; Weissing, 1996; Charlesworth, 1994; Caswell, 2000). This result relies on the assumption that mutants are rare, which allows one to neglect the interactions between individuals carrying the mutant allele in the invasion analysis.

When dispersal is limited due to family or spatial population structure, interactions between mutants can no longer be neglected when evaluating the growth rate; mutant-mutant interactions will occur locally at the level of the interaction group even if the mutant is globally rare. Since the mutant is no longer necessarily locally rare, one needs to track groups with different numbers of mutant alleles (i.e., the local distribution of mutants). In this case, the growth rate *ρ* becomes the eigenvalue of a matrix describing the transitions between different group states (Motro, 1982; Bulmer, 1986; van Baalen and Rand, 1998; Wild, 2011). In this case, the interpretation of the growth rate in terms of individual-centered fitness components is no longer straightforward. In order to understand exactly what the growth rate represents biologically, it needs to be unpacked and expressed in terms of individual-centered properties. Until now, no general interpretation of the mutant growth rate has been provided for group structured populations subject to local heterogeneities, such as demographic or environmental fluctuations.

In the absence of a general and clear interpretation of the growht rate of a mutant allele, several different measures of *invasion fitness*, defined as any quantity allowing to determine the fate of a mutant, have been proposed. One approach computes invasion fitness as the *basic reproductive number, R*_0_, of a mutant lineage (Massol et al., 2009). This gives the total number of successful emigrants produced by a mutant lineage over its lifetime when the lineage was started in a single group by some *distribution of emigrants*. It is well established in mathematical biology that maximizing the basic reproductive number *R*_0_ (the eigenvalue of the next generation matrix associated with the process) is equivalent to maximizing its growth rate (holding the resident population constant), and thus predicts the direction of selection in the same way (Caswell, 2000; Ellner and Rees, 2006).

A closely related approach puts forward the total number of successful emigrants produced by a mutant lineage over its lifetime in a single group that was founded by a *single emigrant*, called *R*_m_, as the appropriate measure of invasion fitness (Metz and Gyllenberg, 2001; Cadet et al., 2003). By assumption, this requires that individuals disperse independently and not in clusters, which excludes propagule dispersal. However, a fitness measure should in general be able to account for propagule dispersal, which is important for understanding the life cycle of many species. This raises the question of the general connection between *R*_0_ and *R*_m_ and their interpretation in terms of individual-centered fitness components.

Further, invasion fitness can also be computed as the personal fitness of a randomly sampled carrier of the mutant allele from the founding lineage (Day, 2001; Lehmann et al., 2015; Mullon et al., 2016), which we refer as *lineage fitness*. In contrast to *R*_0_ and *R*_m_, lineage fitness is expressed in terms of individual-centered fitness components, but it has not yet been generalized to subdivided populations with local heterogeneities.

Among all alternative methods for studying evolution in structured populations, the most popular one, however, has perhaps been the direct fitness method of social evolution theory (e.g., Taylor and Frank, 1996; Frank, 1998; Rousset, 2004; Wenseleers et al., 2010). This approach quantifies the effect on selection of local interactions between individuals carrying a mutant allele by using relatedness coefficients and ascertains the direction of selection on a mutant lineage by way of the *inclusive fitness effect*. The inclusive fitness effect is a weak selection decomposition of the change in the personal fitness of a randomly sampled carrier of the mutant allele into direct effects, resulting from an individual expressing the mutant instead of the resident allele, and indirect effects weighted by relatedness among group members, resulting from group neighbours expressing the mutant. The inclusive fitness effect has helped understand the selection pressure on very diverse phenotypes including the sex-ratio, reproductive effort, genomic imprinting, dispersal, menopause, parasite virulence, interactive behavior, senescence, and niche construction in groups structured populations (e.g., Taylor, 1988; Haig, 1997; Frank, 1998; Gandon, 1999; Taylor and Irwin, 2000; Pen, 2000; Lehmann, 2008; Wild et al., 2009; Lion and Gandon, 2009; Johnstone and Cant, 2010; Ronce and Promislow, 2010; Akçay and Van Cleve, 2012; Lion, 2013).

Despite their apparent differences, inclusive fitness, lineage fitness, or, more generally invasion fitness measures, are in fact tightly connected (Akçay and Van Cleve, 2016). For example, under constant demography, the inclusive fitness effect amounts to evaluating the sensitivity of the number of emigrants *R*_m_ or the growth rate *ρ* with respect to variation in continuous trait values and lineage fitness is equal to *ρ* (Ajar, 2003; Lehmann et al., 2015; Mullon et al., 2016), but the general connection between mutant growth rates, inclusive fitness, lineage fitness, and the reproductive numbers, has not been worked out under arbitrary mutant trait types and selection strength with local demographic and/or environmental heterogeneities.

The aim of this paper is to fill these gaps by providing a general interpretation of the mutant growth rate in terms of individual-centered fitness components and connecting formally to each other the different invasion fitness measures. Our results highlight the conceptual unity underlying invasion fitness and resolve some long standing about how inclusive fitness fits in under arbitrary mutant type and strength of selection.

## Model

### Life-cycle

We consider a population of haploid individuals divided into an infinite number of groups. The population is censused at discrete time demographic periods. In each period, each group, independently from each other, can be in one of a countable number of demographic-environmental states. A state can determine the number of individuals in a group (“demographic” state) and/or any environmental factor affecting all individuals within a group (“environmental” state). Local state fluctuations in the population due to demographic or environmental processes can result in population level patterns of temporal and spatial heterogeneity.

Dispersal may occur between groups by individuals alone or by groups of individuals (i.e., propagule dispersal), but dispersal is always assumed to be uniform between groups in the population; in other words, we consider an island model of dispersal (Wright, 1931). The model allows us to represent classical metapopulation processes with variable local group sizes (Chesson, 1981; Rousset and Ronce, 2004), insect colony dynamics with endogenous growth (Avila and Fromhage, 2015), as well as compartmentalized replication as occurs in the stochastic corrector model for prebiotic evolution (Szathmary and Demeter, 1987; Grey et al., 1995).

We assume that only two alleles can segregate in the population, a mutant allele with type *τ* and a resident allele of type *θ* where the set of all possible types is called Θ. Suppose that initially the population is monomorphic or fixed for the resident type *θ* and that a single individual mutates to type *τ*. Will the mutant “invade” the population and increase in frequency? If the resident type *θ* is such that any mutant type *τ*∈ Θ goes extinct with probability one, we will say that *θ* is *uninvadable*. A state that is uninvadable is an evolutionarily stable state. Our aim is to characterize uninvadability mathematically and biologically.

### The resident demographic equilibrium

Following standard assumptions for the dynamics of mutant-resident substitutions (Eshel and Feldman, 1984; Eshel, 1996; Hammerstein, 1996; Weissing, 1996; Metz et al., 1996), we assume that a mutant can only arise in a resident population that is at its demographic equilibrium, and we start by characterizing this equilibrium. Our main assumption is that the stochastic process describing the state dynamics of a focal group in the resident population is given by a discrete time Markov chain on a countable state space (Karlin and Taylor, 1975; Iosifescu, 2007), where the time scale is that of a demographic period (i.e., the scale at which births, deaths, dispersal, and other events occur).

Because groups may affect each other demographically through dispersal, the transition probabilities for this Markov chain may depend endogenously on the resident population dynamics. But since there is an infinite number of groups, the set of infinite interacting Markov chains (one for each group) can be described as a single (inhomogeneous) Markov chain, whose transition probabilities are functions of the expected value of the process (Chesson, 1981, 1984). We assume that this Markov chain is regular, irreducible and aperiodic (Karlin and Taylor, 1975; Iosifescu, 2007), and thus has a stationary distribution (see Appendix A).

### The mutant multitype branching process

We now introduce a mutant into the backdrop of the resident population in its stationary demographic regime. Denote by *M_t_(s, i)* the random number of groups in the population that are in state *s ∈* 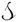 and have exactly *i ∈ I(s)* = {1,2,…,*n*(*s*)} mutant individuals at time *t* = 0,1,2, … where *n(s)* is the number of individuals in a group in state *s* and *t* = 0 is the time of appearance of the mutant. Denote by **M_t_** = 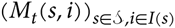 the vector collecting the *M_t_(s, i)* random variables and *e_s_* a vector of the same dimension but whose (*s*, 1)-th component is equal to one, otherwise zero. Starting with a single initial mutant of type *τ* in a focal group in state *s* at time *t* = 0, namely **M**_0_ = **e**(*s*), we are interested in finding a necessary and sufficient condition for the mutant type *τ* to go extinct in finite time with probability one for any state *s ∈* 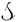 (formally, a condition for Pr [**M**_t_ = **0** for some *t ∈* ℕ | **M**_0_ = **e**(*s*)] = 1 for all *s ∈* 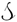).

Since we are interested only in characterizing extinction of the mutant, we assume that it will always remain rare in the total population and approximate the mutant stochastic process as a multitype branching process (Harris, 1963; Karlin and Taylor, 1975; Wild, 2011). It is then sufficient to focus on the (regular) matrix **A** whose (*s′, i′; s, i*) element, denoted *a(s′, i′* | *s, i*), is the expected number of groups in state (*s′, i′*) that are “produced” over one demographic time period by a focal group in state (*s, i*) when the population is otherwise monomorphic for *τ*. It is useful to decompose this as

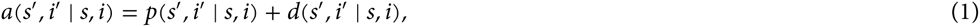

which consists of two terms representing two distinct biological processes. The first is the intra-group (or intra-compartmental) change described by the transition probability *p*(*s′, i′* | *s, i*) that a focal group in state (*s, i*) turns into a group in state (*s′, i′*) after one demographic time period. The second process is the success of a group in replacing other groups by reproduction or fission, which is represented by *d*(*s′, i′* | *s, i*) that measures the expected number of groups in state (*s′, i′*) produced by emigration from, or fission, of a focal group of state (*s, i*). By “producing” a group of state (*s′, i′*), we mean that for a metapopulation process a focal group in state (*s, i*) in a parental generation leaves *i′* ∈ *I*(*s′*) mutant offspring in a group that will be in state *s′* after one demographic time period. For compartmental replication processes (e.g., Grey et al., 1995) this means producing a group in state (*s′, i′*).

### Invasion fitness

It follows from standard results on multitype branching processes (Harris, 1963; Karlin and Taylor, 1975) that the lineage descending from a single mutant *τ* arising in any of the demographic state of the resident *θ* population, will go extinct with probability one if the leading eigenvalue *ρ(τ,θ)* of **Α**(*τ, θ*) is less than or equal to 1. Namely, extinction with probability one occurs if and only if

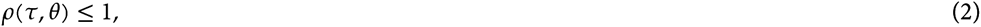

where *ρ* satisfies

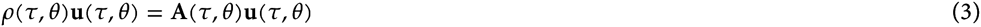

and **u**(*τ, θ*) is the leading right eigenvector of **Α**(*τ, θ*).

The interpretation of *ρ(τ, θ)* is that it gives the asymptotic growth rate of an average trajectory of a mutant lineage; that is, the collection of individuals descending from an individual in which the mutation appeared (Cohen, 1979; Tuljapurkar et al., 2003). In the long-run, the average mutant lineage grows in the direction of **u**(*τ, θ*) so that this vector can be interpreted as a quasi-stationary distribution of group genetic-demographic-environmental states containing at least one individual belonging to the mutant lineage. Namely, element (*s, i*) of **u**, that is *u(s, i)*, is the asymptotic frequency of s-type groups with *i* ≥ 1 mutants; this interpretation holds whether the mutant lineage goes extinct or invades the population (Harris, 1963).

It follows directly from the construction of the model that *ρ(θ, θ)* = 1; namely, the growth of a resident lineage in a resident population is equal to one (see Appendix A for a proof). This implies that a resident type Θ ∈ Θ is uninvadable if, and only if,

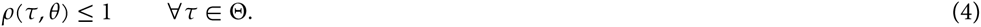

Thus *θ* is uninvadable only if *θ* solves the maximization problem max_τ∈Θ_ *ρ(τ, θ)*.

Now that we have a mathematical characterization of uninvadability in terms of the growth rate *ρ(τ, θ)* of the mutant lineage, we present five different measures of invasion fitness that are all related to *ρ(τ, θ)* and are all expressed in term of biological quantities that have appeared previously in the literature. All these quantities are derived in the Appendix from the elements *a(s′, i′ | s, i), p(s′, i′ | s, i)*, and/or *d(s′, i′ | s, i)* (eq. (1)), and the explicit mathematical expressions are given in Table 1.

**Table 1:**
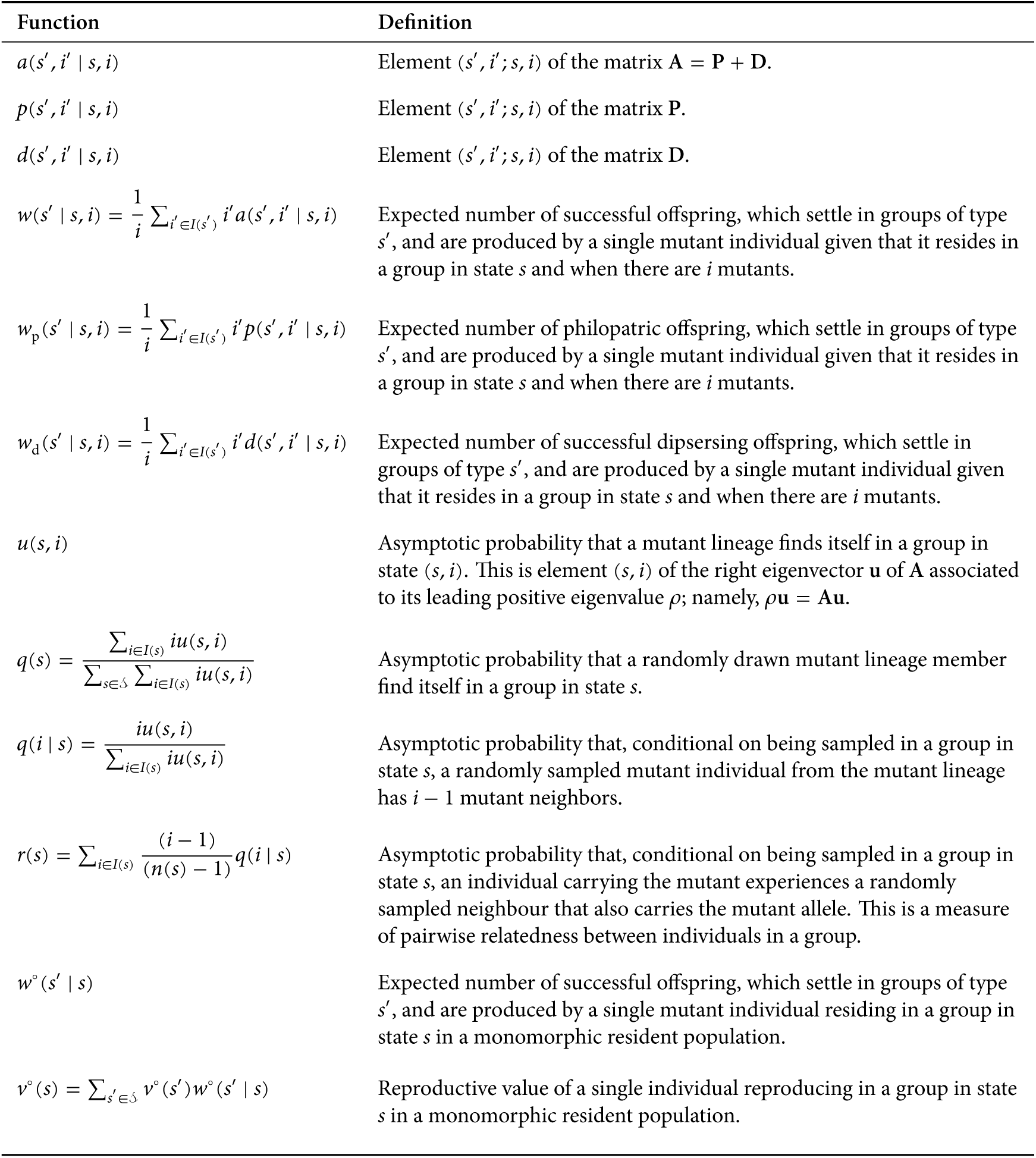
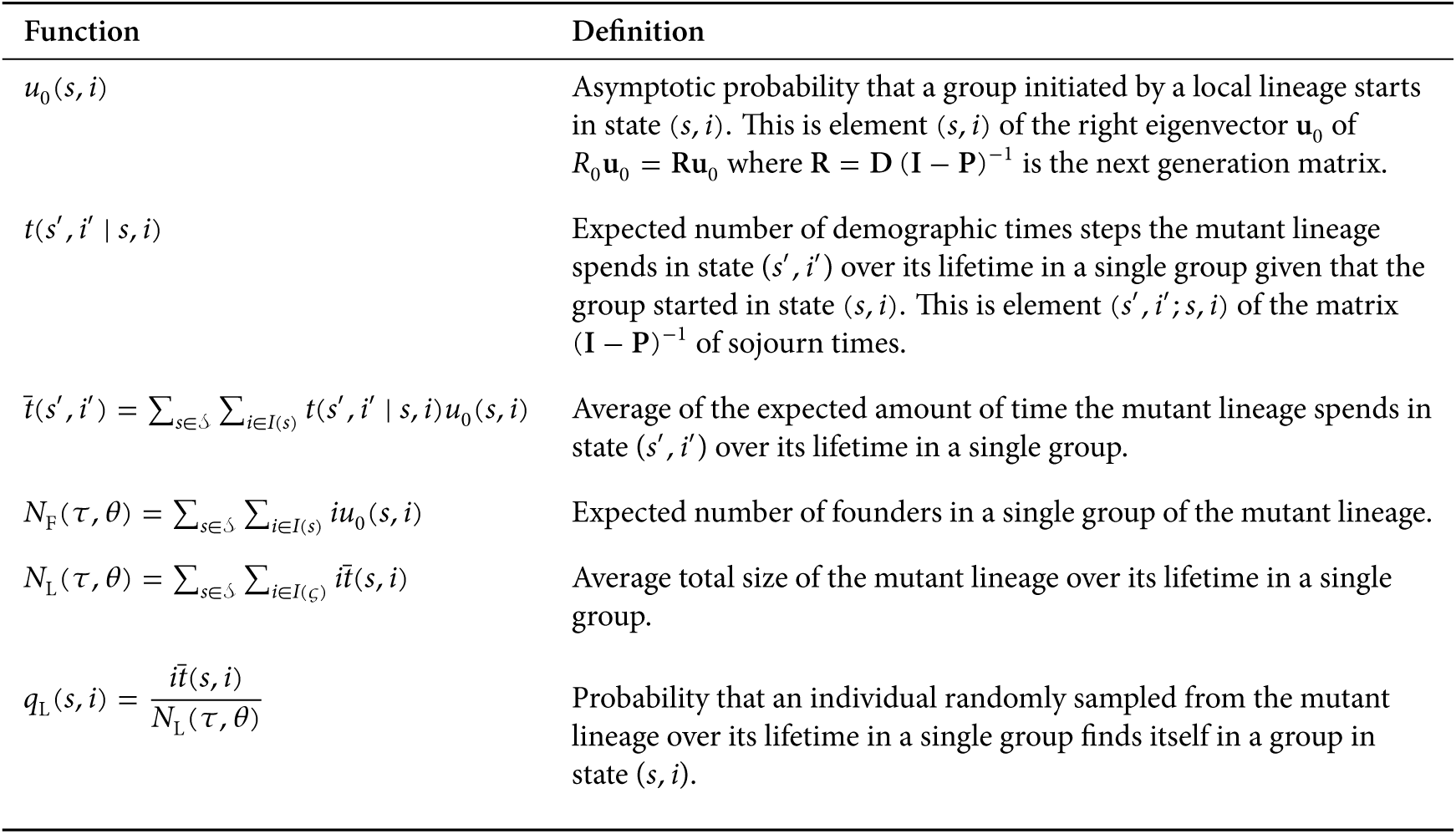
Definitions of the functions and vectors used for lineage fitness, inclusive fitness, and the reproductive number.

| Function | Definition |
| --- | --- |
| $a(s', i' \vert s, i)$ | Element $(s', i'; s, i)$ of the matrix $\mathbf{A} = \mathbf{P} + \mathbf{D}$. |
| $p(s', i' \vert s, i)$ | Element $(s', i'; s, i)$ of the matrix $\mathbf{P}$. |
| $d(s', i' \vert s, i)$ | Element $(s', i'; s, i)$ of the matrix $\mathbf{D}$. |
| $w(s' \vert s, i) = \frac{1}{i} \sum_{i' \in I(s')} i' a(s', i' \vert s, i)$ | Expected number of successful offspring, which settle in groups of type $s'$, and are produced by a single mutant individual given that it resides in a group in state $s$ and when there are $i$ mutants. |
| $w_p(s' \vert s, i) = \frac{1}{i} \sum_{i' \in I(s')} i' p(s', i' \vert s, i)$ | Expected number of philopatric offspring, which settle in groups of type $s'$, and are produced by a single mutant individual given that it resides in a group in state $s$ and when there are $i$ mutants. |
| $w_d(s' \vert s, i) = \frac{1}{i} \sum_{i' \in I(s')} i' d(s', i' \vert s, i)$ | Expected number of successful dispersing offspring, which settle in groups of type $s'$, and are produced by a single mutant individual given that it resides in a group in state $s$ and when there are $i$ mutants. |
| $u(s, i)$ | Asymptotic probability that a mutant lineage finds itself in a group in state $(s, i)$. This is element $(s, i)$ of the right eigenvector $\mathbf{u}$ of $\mathbf{A}$ associated to its leading positive eigenvalue $\rho$; namely, $\rho \mathbf{u} = \mathbf{A} \mathbf{u}$. |
| $q(s) = \frac{\sum_{i \in I(s)} i u(s, i)}{\sum_{s \in \mathcal{S}} \sum_{i \in I(s)} i u(s, i)}$ | Asymptotic probability that a randomly drawn mutant lineage member find itself in a group in state $s$. |
| $q(i \vert s) = \frac{i u(s, i)}{\sum_{i \in I(s)} i u(s, i)}$ | Asymptotic probability that, conditional on being sampled in a group in state $s$, a randomly sampled mutant individual from the mutant lineage has $i - 1$ mutant neighbors. |
| $r(s) = \sum_{i \in I(s)} \frac{(i - 1)}{(n(s) - 1)} q(i \vert s)$ | Asymptotic probability that, conditional on being sampled in a group in state $s$, an individual carrying the mutant experiences a randomly sampled neighbour that also carries the mutant allele. This is a measure of pairwise relatedness between individuals in a group. |
| $w^\circ(s' \vert s)$ | Expected number of successful offspring, which settle in groups of type $s'$, and are produced by a single mutant individual residing in a group in state $s$ in a monomorphic resident population. |
| $v^\circ(s) = \sum_{s' \in \mathcal{S}} v^\circ(s') w^\circ(s' \vert s)$ | Reproductive value of a single individual reproducing in a group in state $s$ in a monomorphic resident population. |
| $u_0(s, i)$ | Asymptotic probability that a group initiated by a local lineage starts in state $(s, i)$. This is element $(s, i)$ of the right eigenvector $\mathbf{u}_0$ of $R_0 \mathbf{u}_0 = R \mathbf{u}_0$ where $R = D(I - P)^{-1}$ is the next generation matrix. |
| $t(s', i' \vert s, i)$ | Expected number of demographic times steps the mutant lineage spends in state $(s', i')$ over its lifetime in a single group given that the group started in state $(s, i)$. This is element $(s', i'; s, i)$ of the matrix $(I - P)^{-1}$ of sojourn times. |
| $\bar{t}(s', i') = \sum_{s \in \mathcal{S}} \sum_{i \in I(s)} t(s', i' \vert s, i) u_0(s, i)$ | Average of the expected amount of time the mutant lineage spends in state $(s', i')$ over its lifetime in a single group. |
| $N_F(\tau, \theta) = \sum_{s \in \mathcal{S}} \sum_{i \in I(s)} i u_0(s, i)$ | Expected number of founders in a single group of the mutant lineage. |
| $N_L(\tau, \theta) = \sum_{s \in \mathcal{S}} \sum_{i \in I(s)} i \bar{t}(s, i)$ | Average total size of the mutant lineage over its lifetime in a single group. |
| $q_L(s, i) = \frac{i \bar{t}(s, i)}{N_L(\tau, \theta)}$ | Probability that an individual randomly sampled from the mutant lineage over its lifetime in a single group finds itself in a group in state $(s, i)$. |

## An ecstasy in five fits: five invasion fitness measures

### Lineage fitness

First, we let the *lineage fitness* of a mutant type *τ* in a resident *θ* population be

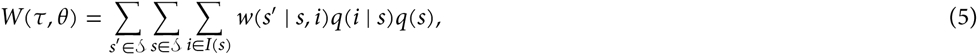

where *w(s′ | s, i)* is the expected number of successful offspring, which settle in groups of type *s′*, given that the parent is a mutant residing in a group in state (*s, i*). Lineage fitness also depends on the probability *q(i | s)* that, conditional on being sampled in a group in state *s*, a randomly sampled mutant individual from the mutant lineage has *i*– 1 mutant neighbors. This can be thought as the conditional mutant *experienced profile distribution* in the stationary mutant distribution, and *q(s)* is the probability that a randomly sampled individual from the mutant lineage finds itself in a group in state s. When there is only one demographic state, *W(τ, θ)* reduces to eq. (A.1) of Day (2001) and eq. (A.7) of Mullon et al. (2016).

Lineage fitness *W(τ, θ)* is the fitness of a randomly sampled carrier of the mutant allele from its lineage, where *w*(*s′ | s, i*) is an individual-centered fitness component variously called “direct”, “personal”, or “individual” fitness in social evolutionary theory (e.g, Frank, 1998; Rousset, 2004), and will be here refered to it as personal fitness. It involves offspring reaching adulthood in the group of the parent and in other groups through dispersal, and can thus also be written as

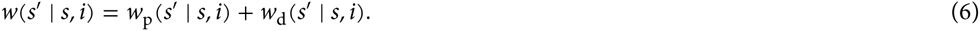

Here, *w_p_* (*s′* | *s, i*) is the expected number of philopatric offspring, which settle in a group in state *s′*, given that the parent is a mutant that reproduced in a group in state (*s, i*), while *w_d_*(*s′* | *s, i*) is such offspring produced by dispersal, and thus reach adulthood in other groups in state s′. This decomposition of personal fitness matches the decomposition of the element of the transition matrix of the mutant given in eq. (1) (see Table 1 and Appendix E, where we further decompose these terms into sub-components that have appeared previously in the literature).

In Appendix B, we show that the growth rate of the mutant lineage is exactly equal to lineage fitness of the mutant; namely,

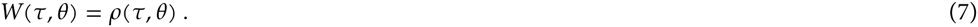

This equation immediately implies that *τ* is uninvadable if it solves the optimization problem max_τ∈Θ_ *W(τ, θ)*. In other words, the type is uninvadable if it “maximizes” lineage fitness. Since lineage fitness is the statistical average over all genetic demographic-environmental states of the personal fitness of the carrier of the mutant allele, it can be interpreted as a gene-centered measure of fitness^1^, since it is the maximand of the number of mutant replica copies produced by a representative individual carrying the mutant allele. The condition for uninvadability (eq. 7) can also be interpreted as a version for class structured population of the seminal uninvadability condition obtained for multilocus systems in panmictic populations, where the statistical average is over multilocus genetic states (Eshel and Feldman, 1984, eq. 10, Eshel et al., 1998, eq. 7).

### Inclusive fitness

Let us now decompose personal fitness as

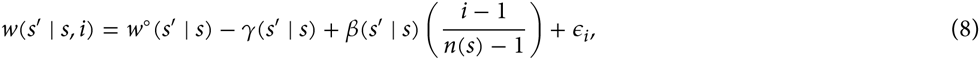

where *w*° (*s′* | *s*) is the expected number of successful offspring, which settle in groups of type *s′*, given that the parent is a resident reproducing in a group in state *s* in a monomorphic resident population, and where the superscript ∘ will throughout denote a quantity that is evaluated in the absence of natural selection, i.e., neutral process determined by the monomorphic resident population. Personal fitness also depends on *γ(s′ | s)*, which is the additive effect on the personal fitness of an individual stemming from it switching to the expression of the mutant allele, *β(s′ | s)*, which is the additive effect on the personal fitness of a mutant stemming from a neighbor switching to the expression of the mutant, and (*i* – 1)/(*n*(*s*) – 1),which is the frequency of mutants in a the neighborhood of a mutant individual in a group with *i* mutants. The *direct* effect *γ (s′ | s)* and the *indirect* effect *β(s′ | s)* are obtained by minimizing the mean squared error *ε_i_* in the linear prediction of personal fitness (see Box 1 for details).

#### Box I Weighted least square regression

We here show how to obtain the cost *γ*(*s′* | *s*) and benefit β(*s′* | *s*) in eq. (8). These are found by minimizing for each state *s* ∈ 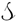 the sum of squared errors *ε_i_* weighted by the probabilities *q*(*i* | *s*):

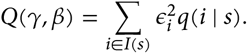

That is, from eq. (8), we minimize

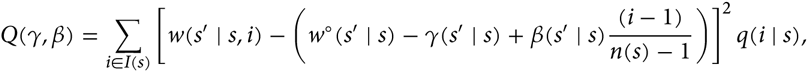

with respect to *γ* and *β* From the prediction theorem for minimum square error prediction (Karlin and Taylor, 1975, p. 465), we then have 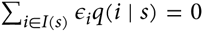 for all *s* ∈ 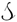 which is one of the main feature we use to obtain the expression for inclusive fitness (see Appendix C).

We let the *inclusive fitness* of a mutant type *τ* in a population with residents of type *θ* be

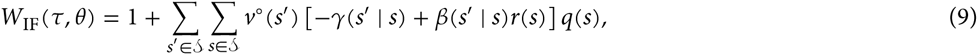

where *v*° (*s*) is the *neutral reproductive value* of a single individual reproducing in a group in state *s*. This is the relative asymptotic contribution of an individual in state *s* to the population (see Taylor, 1996 and Rousset, 2004 for an introduction to this concept). Inclusive fitness also depends on the probability *r(s)* that, conditional on being sampled in a group in state *s*, an individual carrying the mutant experiences a randomly sampled neighbour that also carries the mutant allele. This is a measure of pairwise relatedness between two individuals in a group (see Table 1). In a monomorphic resident population, relatedness [then given by *r*° (*s*)] reduces to the standard concept of probability of identity by descent between two randomly sampled group members (e.g., Frank, 1998; Rousset, 2004). In sum, the inclusive fitness *W*_IF_(*τ,Θ*) of a randomly sampled mutant from the lineage distribution *q(s)* is the reproductive-value weighted average personal fitness cost *γ (s′ | s)* of carrying the mutant allele and the relatedness weighted personal indirect fitness benefit *β(s′ | s)* of carrying the mutant.

We show in Appendix C that inclusive fitness *W*_IF_(*τ, θ*) predicts whether or not the mutant invades in

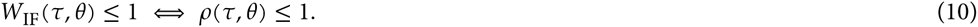

Hence, a strategy is uninvadable if and only if inclusive fitness is maximized, in the sense that *τ* solves the problem max_τ∈Θ_ *W*_IF_ (*τ, θ*). This shows that, regardless of the force of selection, uninvadability can be expressed in terms of the three standard measures of “value” emphasized by social evolution theory (Frank, 1998): (i) the direct cost and indirect benefit within each class of an individual expressing the mutant, (ii) the pairwise relatedness between interacting individuals, and (iii) the neutral reproductive value of the descendants in each class. It is important to note that the inclusive fitness *W*_IF_(*τ,θ*) is not equal to the growth rate *ρ(τ, θ)*, but is a linear function of it (see eq. (C.5) in Appendix C).

### Reproductive numbers

We let the *basic reproductive* number of a mutant type *τ* in a resident *θ* population be

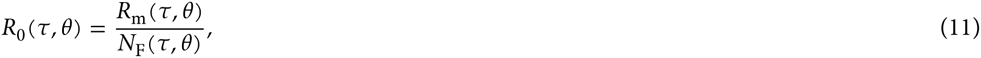

which depends on the expected number *N*_F_(*τ, θ*) of mutant colonizing the same group and descending from the same natal group, and on the successful number of emigrants

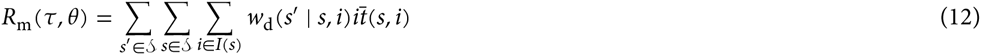

produced by all individuals of the mutant lineage over its lifetime in a single group. This depends on the expected number *w*_d_(*s′* | *s, i*) of emigrant offspring that settle in groups of type *s′* (see eq. (6)) and on the total expected amount of time 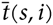 that a mutant lineage spends in a single group in state (*s, i*) in the asymptotic distribution of the mutant lineage. In sum, the basic reproductive number gives the expected number of successful emigrants produced by a lineage during its whole sojourn time in a single group and until its local extinction in that group, relative to the expected number of founders of such a lineage. Although the expression on the right-hand of eq. (12) does not appear previously in the literature, it precisely corresponds to the mathematical definition of the basic reproductive number given in the literature (Caswell, 2000; Ellner and Rees, 2006, see Appendix D). Further, when there is only one demographic state, *R*_m_(*τ, θ*) reduces to eq. (3) of Ajar (2003).

In Appendix D, we show that the basic reproductive number *R*_0_ (*τ, θ*) predicts whether or not the mutant invades in the same way as the growth rate *ρ(τ, θ);* namely

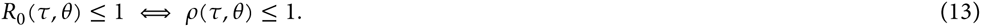

Hence, a strategy is uninvadable if the basic reproductive number is maximized. Suppose now that the number of founders *N*_F_(*τ, θ*) is independent of the mutant; an example would be *N*_F_ (*τ, θ*) = 1 so there can be no propagule dispersal and individuals can only migrate independently of each others. Then, uninvadability can be characterized in terms of *R_m_(τ, θ)* alone:

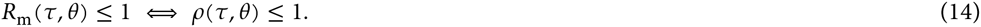

Hence, a strategy is uninvadable if the expected number *R_m_(τ, θ)* of successful emigrants is maximized.

Both reproductive numbers, *R*_0_ and *R*_m_, count (emigrant) successful offspring as produced by a whole set of individuals in the lineage, and, by contrast to *W*(*τ, θ*) and *W*_IF_(*τ, θ*), are thus not individual-centered. In order to have a reproductive number that is expressed in terms of the personal fitness of a representative carrier of the mutant, we let the *lineage fitness proxy* of a mutant type *τ* in a resident *θ* population be given by

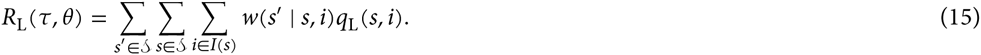

Here, *q*_L_ (*s, i*) is the probability that an individual randomly sampled from the mutant lineage over its lifetime in a single group finds itself in a group in state (*s, i*) (see Table 1). This expression is a direct analogue to lineage fitness, with the only difference that the probability distribution *q*_L_ (*s, i*) depends on the lifetime of the lineage in a single group, and not on the asymptotic lineage distribution *u(s, i)* as does lineage fitness. When there is only one demographic state, *R*_L_(*τ, θ*) reduces to eq. (3) of Lehmann et al. (2015).

We show in Appendix D that lineage fitness proxy *R*_L_(*τ, θ*) predicts whether or not the mutant invades in the same way as the growth rate *ρ(τ, θ);* that is,

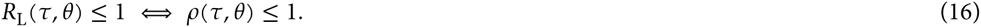

An uninvadable strategy thus also maximizes lineage fitness proxy.

### Results summary

Summarizing all the above results, we have shown that the growth rate is equal to lineage fitness *ρ(τ, θ)* = *W*(*τ, θ*) and the following characterizations of the condition under which a mutant goes extinct are equivalent:

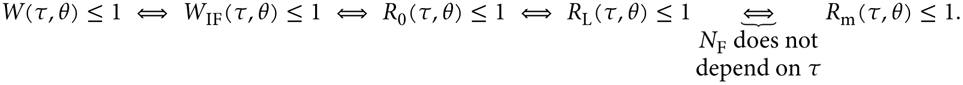

## Discussion

Our results show that the different invasion fitness measures that have been proposed so far all equivalently determine which strategy is uninvadable, and that they can all be connected through their relationship to the growth rate of a mutant allele. The mathematical theory we present provides a formal framework for understanding the broad notion that different fitness measures must align (e.g., Metz et al., 1992; Roff, 2008; Akçay and Van Cleve, 2016). Our results also reveal interesting features of the different invasion fitness measures, which we now discuss.

### Lineage and inclusive fitness

Uninvadability can be equivalently characterized in terms of lineage fitness or inclusive fitness. This duality is interesting as these two gene-centered invasion fitness measures are expressed in terms of different individual-centered fitness components experienced by representative carriers of the mutant allele. Lineage fitness is expressed only in terms of the personal fitness of a randomly drawn individual carrying the mutant allele, where the carrier is drawn from the distribution of group states experienced by members of the mutant lineage (all genetic-demographic-environmental states). In contrast, inclusive fitness is expressed in terms of the direct fitness cost and relatedness weighted indirect fitness benefit accruing to a randomly drawn carrier of the mutant allele. Writing fitness in terms of cost and benefit requires making a comparison between the number of offspring produced by an individual expressing the mutant allele relative to expressing the resident allele. But in order for this comparison to be unbiased, how the fitness value of an offpsring depends on the demographic and/or environmental state in which it settles must be taken into account. Thus, each offspring needs to be appropriately weighted.

Importantly, we find that these weights are the neutral reproductive values of the monomorphic resident population regardless of the strength of selection on the mutant. The intuitive reason for this result is that reproductive value weighting “converts” number of offspring in different states into their proportionate contribution to the population. By choosing the conversion factors to be the neutral reproductive values of the resident allele, the inclusive fitness directly allows determining the increase (or decrease) in descendants into the far future that a typical carrier of the mutant allele leaves relative to the typical carrier of the resident allele, in a monomorphic resident population. This result is consistent with previous population genetic formulations of allele frequency change in class-structured populations under arbitrary strength of selection (Lehmann and Rousset, 2014). Our analysis thus generalizes the exact version of inclusive fitness (e.g., Queller, 1992; Frank, 1997; Gardner et al., 2011) to class-structured populations with variable number of interaction partners, and shows that the standard neutral reproductive value weighting (e.g., Taylor and Frank, 1996; Rousset, 2004) is maintained in this generalization.

Inclusive fitness makes explicit that the force of selection on a mutant allele depends on (i) how individuals in different demographic and environmental states contribute differently to the gene pool and on (ii) the genetic association between individuals due to local common ancestry, regardless of the complexity of the biological situation at hand and the strength of selection. These biological features, hidden in the other invasion fitness measures, also become apparent if one considers only the first-order effects of selection on the growth rate when the evolving traits have continuous values. This is the situation usually considered in the adaptive dynamics and inclusive fitness literature where one looks for evolutionary attractors (Taylor, 1996; Geritz et al., 1998; Rousset, 2004; Dercole and Rinaldi, 2008). In this situation, the sensitivity of the growth rate with respect to changes in trait value boils down to the inclusive fitness effect derived previously by the direct fitness method (Taylor and Frank, 1996; Rousset, 2004, see Box 2 and Appendix E.2 for this connection). Hence, our model makes explicit that the direct fitness method amounts to computing the sensitivity of the growth rate of the mutant with respect to changes in mutant strategy under a general class structure and with environmental heterogeneity (see also Rousset, 2004, pp. 194-196 for a conjecture on that point).

#### Box II Sensitivity of the growth rate

We here provide an expression for the sensitivity of the growth rate when the mutant trait value is varied; that is, the derivative of the growth rate when Θ = ℝ, which is sufficient to evaluate singular strategies and convergence stable states (Taylor, 1996; Rousset, 2004). In Appendix E.2, we prove that the sensitivity of the growth rate is

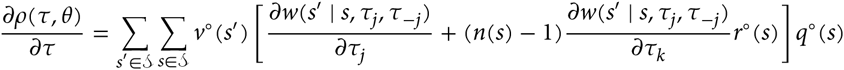

where *w*(*s′* | *s, τ_j_, τ_−j_*) is the personal fitness of an individual with phenotype *τ*_j_, when its group members have phenotype profile *τ_−j_* = (*τ*_1_, …, *τ_j−1_*, … *τ_j+1_*, … *τ_n(s)−1_*), which is the vector collecting the phenotypes of the *n(s)*−1 neighbors of an individual *j* and *k ≠ j*, and all derivatives are evaluated at the resident values Θ. Note that here, both the probability *q*°(*s*)that a mutant experiences a group in state s and relatedness *q*°(*s*) are evaluated in a monomorphic resident population (neutral process). Given further specific biological assumptions on the underlying demographic process, we then recover from the above derivative the expression for the inclusive fitness effect derived by the direct fitness method for the island model (Taylor and Frank, 1996; Rousset and Ronce, 2004, see Appendix E.3.1).

Our analysis thus demonstrates connections between the various theoretical approaches for characterizing adaptations in heterogeneous populations. But depending on the type of questions and insight desired, either inclusive or lineage fitness formulations might be better suited. For instance, lineage fitness may be easier to measure, as it only relies on measuring personal fitness of a representative sample of individuals of the mutant type (see Akçay and Van Cleve, 2016 for further discussions on using invasion fitness measures for empirical system).

### Reproductive numbers

We also derived an explicit expression for the basic reproductive number, *R*_0_, for a group-structured population, which was shown to depend on the ratio of the total lifetime number *R*_m_ of successful emigrants produced by a typical group colonized by members of the mutant lineage, to the expected number *N*_F_ of colonizers of such a typical group. The basic reproductive number is the usual invasion fitness proxy in evolutionary biology and epidemiology (Caswell, 2000; Ellner and Rees, 2006) and is usually used as it simplifies the characterization of the condition under which a mutant invades. It circumvents the need to compute explicitly the growth rate *ρ*, (the eigenvalue of the transition matrix **A**), and only requires a matrix inversion (see Appendix D). When individuals disperse independently and not in clusters (i.e., no propagule dispersal), the basic reproductive number reduces to the number of successful emigrants *R*_m_. Mathematically however, our expression for *R*_m_ (eq. (12)) differs from the expression of *R*_m_ initially introduced as a measure of invasion fitness by Metz and Gyllenberg (2001), insofar as the frequency distribution of the group states of a typically colonized group may depend on the mutant type, which is consistent with the formal proof of *R*_m_ derived by Massol et al. (2009).

Two further points are worth mentioning concerning the reproductive numbers, *R*_0_ and *R*_m_. First, while no relatedness appears explicitly in them, they take inclusive fitness effects into account in the same amount as inclusive fitness (or lineage fitness) does. Second, the reproductive numbers count successful emigrant offspring produced by a whole set of individuals, and thus do not give net successful offspring produced by a representative carrier of the mutant allele. In order to have a fully individual-centered measure of invasion fitness, which keeps the attractive computational features of the reproductive numbers, we derived an expression for lineage fitness proxy *R*_L_ This is the personal fitness of a mutant lineage member randomly sampled from the distribution quantifying the lifetime of the mutant lineage in a local group. This allows one to determine uninvadability with the same generality and simplicity as *R*_0_, but with the same biological interpretation as lineage fitness.

### Generalizations

To obtain our results, we assumed a population of infinite size but allowed for limited dispersal between any local group and local demographic and/or environmental state. This allows one to describe, in at least a qualitative way, different metapopulation processes as well as group (or propagule) reproduction processes subject to local demographic and environmental stochasticity. Conceptually, our qualitative results concerning the generic form of the fitness measures should carry over to isolation-by-distance models and finite total population size once the growth rate is interpreted as the fixation probability.

We also only considered halploid reproduction, but diploid reproduction would not produce qualitatively different results concerning the expressions of lineage fitness, inclusive fitness, or the reproductive numbers. In the case of diploidy, one needs to add an additional class structure within each demographic state so that individuals are either homozygous or heterozygous and produce these two types of offspring. The same extension is needed for class structure such as age or stage (see Box 3 and Appendix F for an example involving stage structure). An extension to continuous classes is also straightforward as its suffices to replaces eigenvectors by eigenfunctions in the characterization of the growth rate (Harris, 1963), and all other calculations should carry over conceptually unchanged (but replacing sums by appropriate integrals). Our approach, however, breaks down when there are global environmental fluctuations affecting all groups in the population simultaneously, in which case the stochastic growth rate needs to be used to ascertain uninvadability (Svardal et al., 2015). Hence, a completely general interpretation of the growth rate of a mutant in terms of individual-centered fitness components, covering all possible biological heterogeneities, is still lacking. But for local heterogeneities, there is a generality and consistency in the interpretation of the force of a selection on a mutant allele that befits the generality of natural selection.

#### Box III Lineage and inclusive fitness for class-structure under fixed demography

Suppose that each group is of constant size but that each individual within a group can belong to one of *n_c_* classes where the set of classes is *C* = {1, …, *n_c_*}. An example would be age structure due to overlapping generations or different castes of social insects like workers and queens. For such a class structured population, we show in Appendix F that the lineage fitness of a mutant *τ* in a *θ* population is

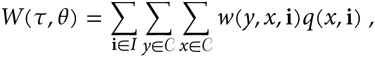

where *w(y, x, i)* is the expected number of class y offspring produced by a class *x* mutant when in a group in state i = (*i*_1_, …, *n_c_*) ∈ *I*, which is the vector of the number of mutant alleles in class 1 to *n_c_*. Here, *I* = (*I*_1_ × … × *I_n_c__*) is the set of possible group states with *I_x_* = {0, 1, …, *n_x_*} being the set of the number of mutant alleles in class *x* and *n_x_* is the number of individuals in that class. In complete analogy with the demographically structured population case, *q*(*x*, i)is the probability that a randomly sampled lineage member finds itself in class *x* and its group state is **i**. The inclusive fitness expression for this model is

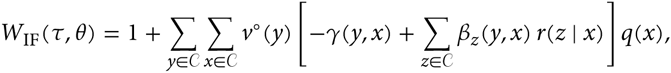

where *q(x)* is the probability that a randomly sampled individual from the mutant lineage finds itself in class *x, γ(y, x)* is the additive effect on the number of class *y* offspring produced by a class *x* individual when expressing the mutant instead of the resident allele, *β_z_(y, x)* is the additive effect on this fitness stemming from group neighbors in class *z* expressing the mutant instead of the resident allele, and *r(z | x)* is the probability that, conditional on being sampled in class x, an individual carrying the mutant experiences a randomly sampled neighbour in class z that also carries the mutant allele.

## Footnotes

1 EA and JVC prefer the nomenclature “gene-lineage-centered”(Akçay and Van Cleve, 2016).

## Appendix A: Properties of the monomorphic resident population

The demographic equilibrium for a monomorphic resident *τ* population described in the main text can be expressed as

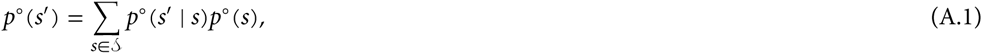

where *p*° (*s*) is the neutral stationary probability that in group is in state *s* and *p*° (*s′* | *s*) denotes the neutral transition probability from state *s* to *s′* (possibly depending endogenously on the distribution *p*° (*s*)).

We now prove that in a monomorphic *θ* population the neutral transition matrix A° has dominant eigenvalue *ρ(θ, θ)* = 1. We do so by constructing a positive left eigenvector v° > 0 of A° with unit eigenvalue (i.e., such that v°A° = v°). Then, since A° is irreducible and non-negative (and v° > 0), the Perron-Frobenius theorem tells us that the dominant eigenvalue of A° is one (e.g., Karlin and Taylor, 1975). We construct v° = (*v*°(1,1), …, *v*°(1,*n*(1)), *v*°(1,2),…) with (*s, i*) element

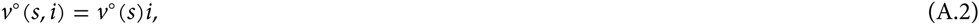

where *v*°(*s*) > 0 corresponds to the reproductive value of an individual in class *s* (see Taylor, 1996 and Rousset, 2004). By definition, reproductive values satisfy

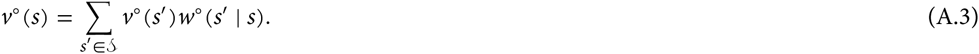

To show that our construction of v° is a left eigenvector of A°, we first write the (*s, i*) element of v°A° by using eq. A.2 as

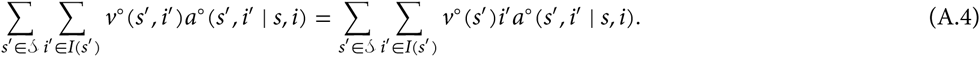

Then, we note that the total expected number of mutant individuals in a group of type *s′* produced by a group of type (*s, i*) can be written in two ways,

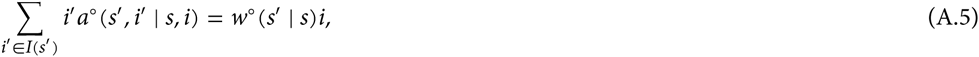

where, owing to neutrality, fitness *w*° (*s′* | *s*) is independent of *i*. Using eq. (A.5) first and (A.3) second, the (*s, i*) element of v°A° can thus be written as

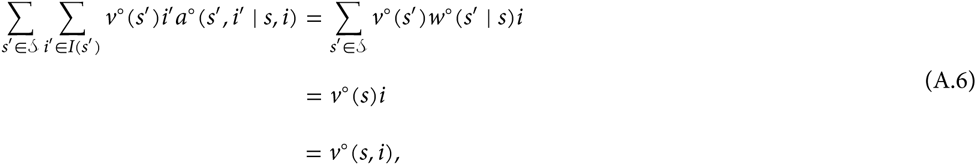

i.e., as the (*s, i*) element of v°, which shows that our construction of v° is indeed a left eigenvector of A° with unit eigenvalue, as required.

## Appendix B: Lineage fitness

We here prove that *ρ*(*τ, θ*) = *W*(*τ, θ*) (eq. (7) of the main text). To that aim, we first note that eq. (A.5) holds out of neutrality and that

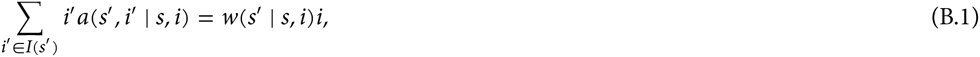

since the right hand side is the total expected number of mutant individuals in a group of type *s′* produced by a group of type (*s, i*). Second, we let n = (1, 2, …, *n*(1), 1, 2 …, *n* (2), …, *n(s)*) and premultiply *ρ**u*** = Au by **n** gives **n** · *ρ***u** = (**n** · **Au**), where · is the dot product. Using eq. (B.1) we have

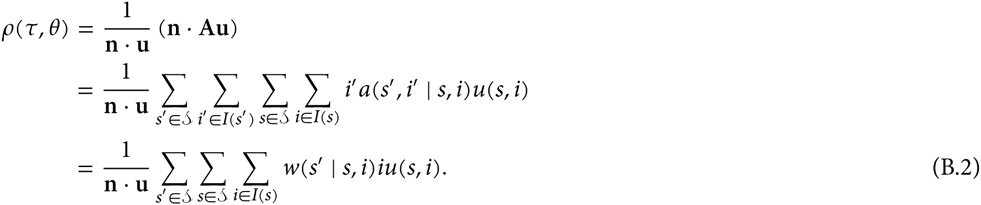

Using the definitions of *q*(*i* | *s*) and *q*(*s*) given in the Table 1 of the main text (where 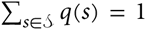 and 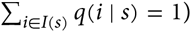 *q*(*i* | *s*) = 1), we can then write

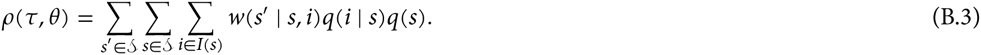

The right hand side is exactly *W*(*τ, θ*), whereby *ρ*(*τ, θ*) = *W*(*τ, θ*).

## Appendix C: Inclusive fitness

Here, we prove that the uninvadability condition can be expressed in terms of inclusive fitness (eq. (10)). For this, we premultiply *ρ***u** = **Au** by v°, which gives v° · *ρ***u** = (**v**° · **Au**). Using eq. (A.2) then entails

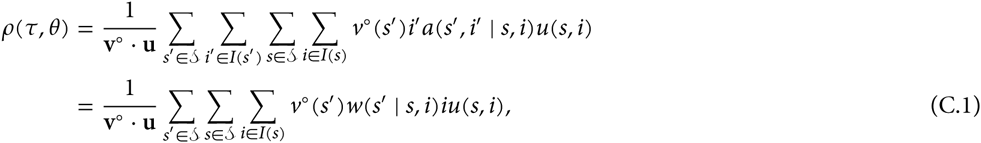

and using

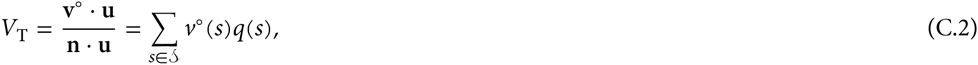

which is the average reproductive value, yields

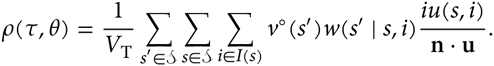

Using

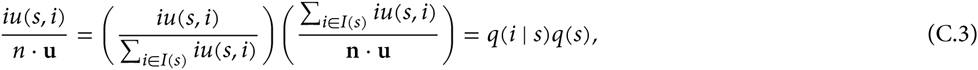

we have

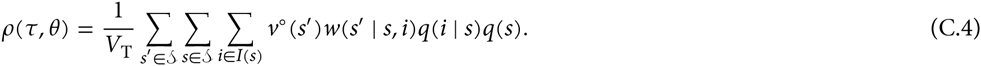

We now use the regression equation form for *w*(*s′* | *s, i*) (eq. (8) of the main text), insert it into eq. (C.4) and obtain

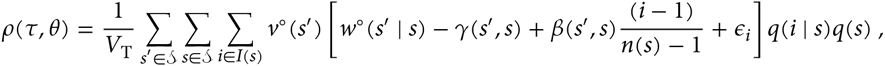

which becomes

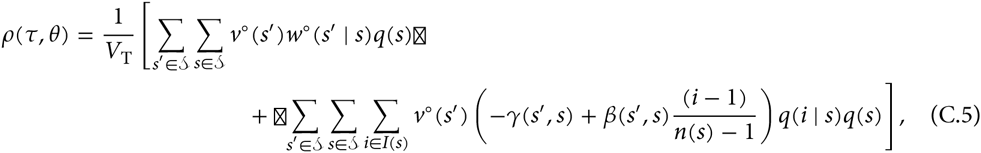

since the minimum mean square error used to obtain *γ*(*s′, s*) and *β*(*s′, s*) ensures that 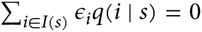 for all *s ∊* 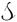 (see Box 1). Using eq. (A.3), the double sum in the first line of eq. (C.5) is seen to be *V*_T_, and using the definition of relatedness *r*(*s*) = Σ_*i∊I(s)*_ [(*i* – 1)/(*n*(*s*) – 1)] *q*(*i* | *s*) (see Table 1), we can simplify the sum on the second line of eq. (C.5) using the expression for inclusive fitness (eq. (9) of the main text) to obtain

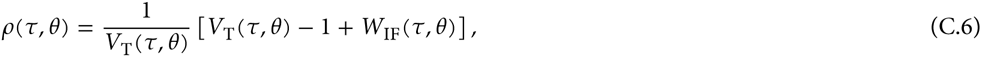

whence

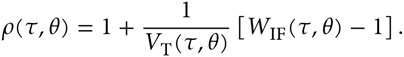

Since, *V*_T_ (*τ, θ*) > 0, we finally have

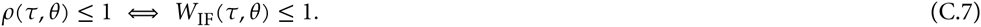

Hence, a type *τ* is uninvadable if it solves max_τ∊Θ_ *W*_IF_ (*τ, θ*).

## Appendix D: Reproductive numbers

### D.1 Basic reproductive number and expected number of emigrants

Here, we prove the uninvadability condition expressed in terms of the basic reproductive number (eq. (13) of the main text and Table). According to our notations, the mean matrix of the branching process can be decomposed as

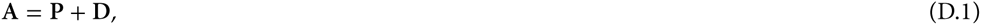

where **P** is the matrix collecting the *p*(*s′, i′ | s, i*) elements and **D** is the matrix collecting the *d(s′, i′ | s, i*) elements (see eq. (1) or Table 1). Then an application of the next generation theorem (Caswell, 2000; Thieme, 2009) shows that

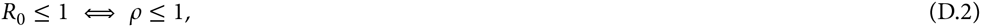

where *R*_0_ is the leading eigenvalue of the next generation matrix

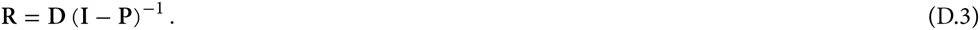

This matrix has leading right eigenvector **u**_0_ whose element *u_0_ (s, i)* is the asymptotic probability that a group initiated by a local lineage starts in state (*s, i*) (*R*_0_ **u**_0_ = **Ru**_0_). The elements of **R** are

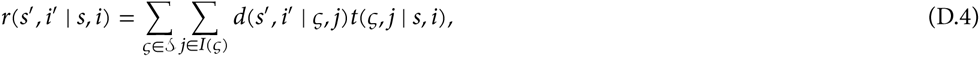

where *t*(*ζ, j | s, i*) is the expected number of demographic time steps the mutant lineage spends in state (*ζ, j*) over its lifetime in a single group given that the group started in state (*s, i*). These sojourn times are elements of the “fundamental matrix” (**I** – **P**)^−1^ (Grinstead and Snell, 1997). The interpretation of *r*(*s′, i′* | *s, i*) is that it gives the total expected number of groups in state (*s′, i′*) produced through dispersal over the lifetime of the mutant lineage in a single group that started in state (*s, i*).

Using the above, we now rewrite *R*_0_ using the same line of argument as for lineage fitness. Hence, we first let

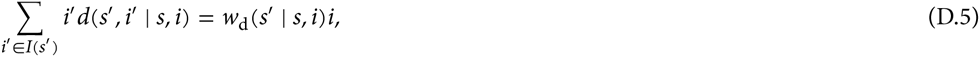

where *w*_d_(*s′* | *s, i*) is the total expected successful number of immigrants in groups in state *s′* produced by a single mutant in a group in state (*s, i*). Premultiplying *R*_0_ **u**_0_ = **Ru**_0_ by **n** and using eq. (D.5) entails that

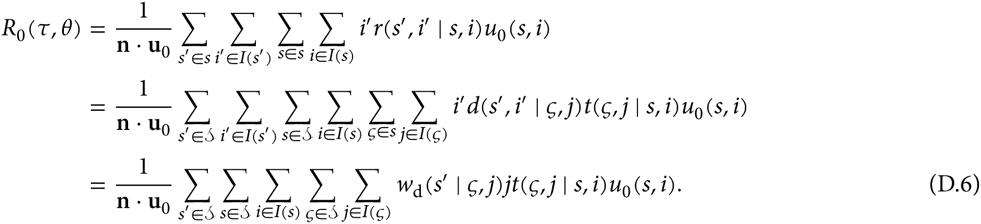

In order to further simplify *R*_0_ we set

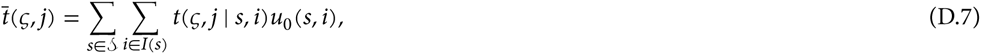

which is the average of the expected amount of time the mutant lineage spends in state (*ζ, j*) over its lifetime in a single group. We also let

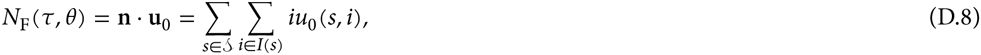

which is the expected number of founders of the mutant lineage. By further denoting

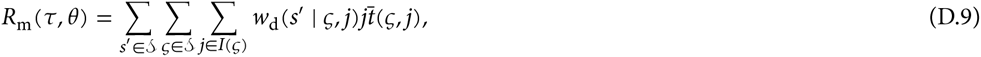

and inserting into eq. (D.6), we have

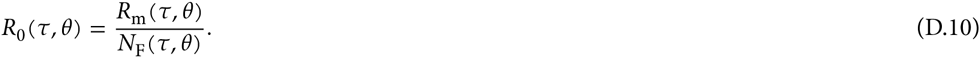

### D.2 Lineage fitness proxy

We will now rewrite eq. (D.10) in terms of lineage fitness proxy (eq. (15) of the main text). For this, we set

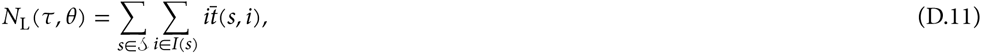

which is the expected total size of the mutant lineage over its lifetime in a single group. Extending the argument of Mullon and Lehmann (2014, Appendix A), this is also

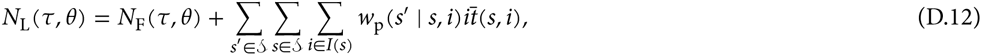

since *N*_F_(*τ, θ*) is the expected number of mutant individuals founding a single group and the sum is the expected number of mutant offspring settling locally and produced over the lifetime of the lineage in that group. Subtracting eq. (D.11) from eq. (D.12), inserting into eq. (D.9) and using eq. (6), we can write

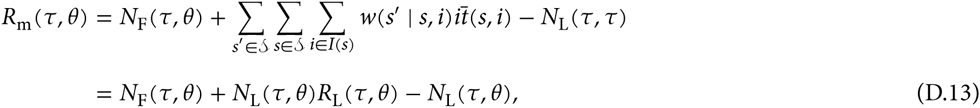

where the second line follows from using eq. (15). Inserting eq. (15) of the main text and eq. (D.13) into eq. (D.10) gives

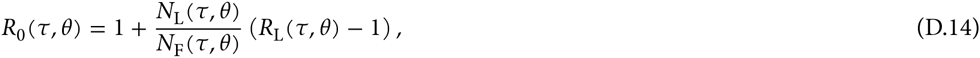

which shows that *R*_0_(*τ, θ*) ≤ 1 ⇔ *R*_L_(*τ, θ*) ≤ 1, whereby

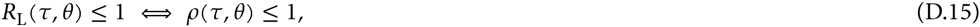

## Appendix E: Connections to previous work

We here provide different connections to fitness components that appear in the literature.

### E.1 Fitness decomposition: philopatric and dispersed

We start by further decomposing the two fitness components in eq. (6). First, we can write

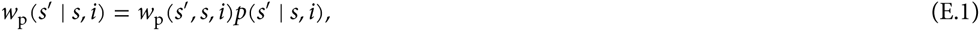

where *p*(*s′* | *s, i*) is the probability that a group will be in state *s′* in the offspring generation given that it was in state (*s, i*) in the parental generation and *w*_p_(*s′, s, i*) is the expected number of successful philopatric offspring given that the offspring settle in a group in state *s′* and the parent reproduces in a group in state (*s, i*). We can also write

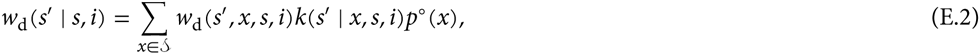

where *p*° (*x*) is the (neutral) probability that a group randomly sampled in the monomorphic resident population is in state *x*. Here, *k*(*s′* | *x, s, i*) is the probability that a group that was in state (*x*, 0) in the parental generation and has been colonized by a mutant descending from a group in state (*s, i*) will become a group in state *s′* in the offspring generation, and *w*_d_(*s′, x, s, i*) is the expected number of dispersing offspring that a single mutant produces given that it resides in a group in state (*s, i*) and given that the group where the offspring settle is in state *s′* in the offspring generation and was in state *x* in the parental generation (with 0 mutants). The conditional fitness functions *w*_p_(*s′, s, i*) and *w*_d_(*s′, x, s, i*) are the elementary individual-based fitness components of models in demographically structured populations (e.g., eqs. 31-32 of Rousset and Ronce, 2004).

We now prove the expressions for the two above conditional expectations (eqs. (E.1)-(E.2)). From Table 1, the first conditional expectation can be written as

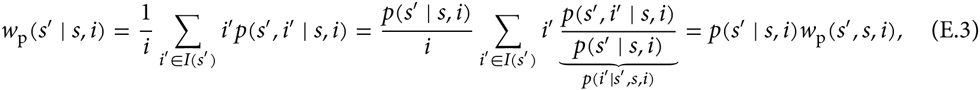

where *p*(*i′* | *s′, s, i*) is the probability that a group will have *i′* mutants in the offspring generation given that it is in state (*s, i*) in the parental generation and in state *s′* in the offspring generation. Here, we used

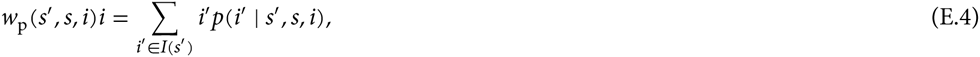

where *w*_p_(*s′, s, i*) the expected number of successful philopatric offspring that a single mutant produces given that it resides in a group in state (*s, i*) and that the group state in the offspring generation is *s*.

From Table 1, the second conditional expectation is

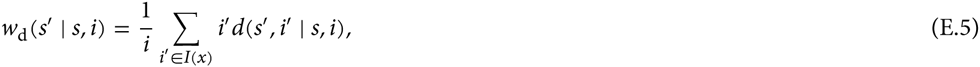

where, conditioning on the state of the group in the parental generation where the offspring disperse to, we can write

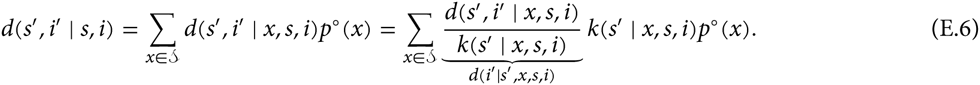

Here, we used in the conditioning the neutral probability *p*° (*x*) that a group randomly sampled in the monomorphic resident population is in demographic state *x*, since dispersing offspring can only land in a group whose state in the parental generation is determined by the resident dynamics. The term *d*(*s′, i′* | *x, s, i*) is the expected number of groups in (*s′, i′*) produced by a group in state (*s′, i′*) and given that they were in state (0, *x*) in the parental generation (with 0 mutants). We now let

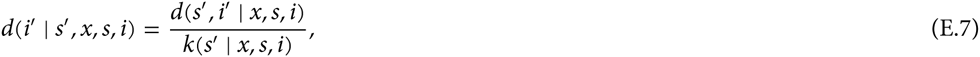

where *k*(*s′* | *x, s, i*) is the probability that a group will be in state *s′* in the offspring generation, given that it was in state (*x*, 0) in the parental generation and has been colonized by a mutant descending from a group in state (*s, i*). Further we have

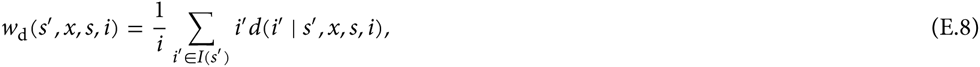

which is the expected number of dispersing offspring that a single mutant produces given that it resides in a group in state (*s, i*) and given that the group where the offspring settle is in demographic state *s′* in the offspring generation and was in state (*x*, 0) in the parental generation. Substituting into eq. (E.6), we then obtain eq. (E.2).

### E.2 Connection to the direct fitness method

We now connect our results to the direct fitness approach (Taylor and Frank, 1996; Rousset, 2004), which, formally, consists of computing the selection gradient on a mutant type when mutant phenotypic deviations are small relative to the resident and is sufficient to evaluate the condition of convergence stability under essentially all conditions (Rousset, 2004; Lehmann and Rousset, 2014). Hence, results from the direct fitness method should match *∂ρ(τ, θ)/∂τ* when the type space is real valued and one dimensional (Θ = ℝ), which we henceforth assume.

### E.3 Sensitivity of the growth rate

To prove the connection to the direct fitness approach we first derive a generic expression for the growth rate sensitivity *∂ρ(τ, θ)/∂τ* under our model assumptions. To that aim, we rewrite the growth rate by using eq. (C.4) as

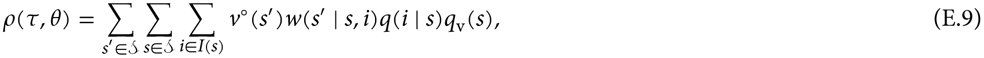

where

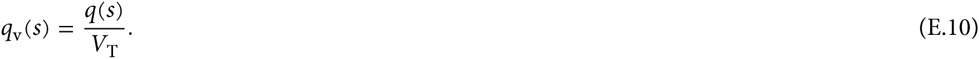

Since, *v*° (*s*′) depends only on the resident, we have

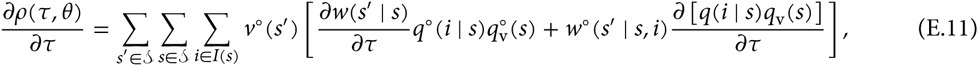

where all derivatives, here and throughout, are evaluated at *τ = θ*. Using the neutral reproductive values (eq. (A.3)), we have

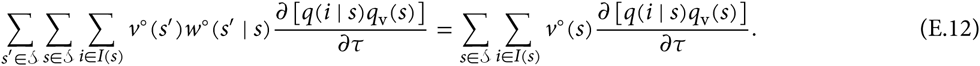

Further, we have

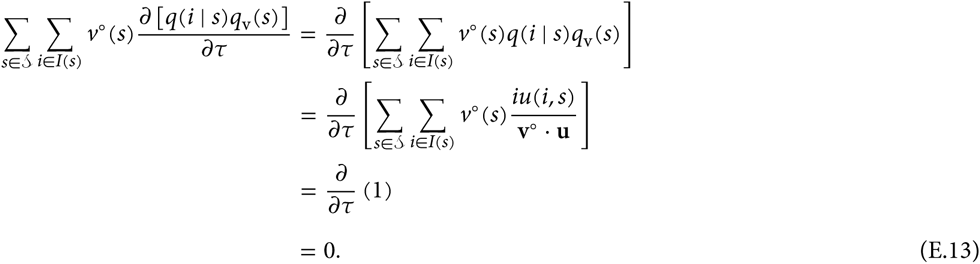

Hence, substituting eq. (E.13) into eq. (E.11) using eq. (E.10) gives

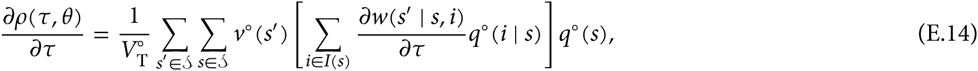

where without loss of generality we can normalize the elements *v°* (*s′*) such that 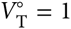.

Note that *w*(*s′* | *s, i*) is the personal fitness of a mutant with phenotype *τ* when its group members consist of *i* – 1 individuals with phenotype *τ* and *n(s)* – *i* individuals with phenotype *θ*. Thus, we can write

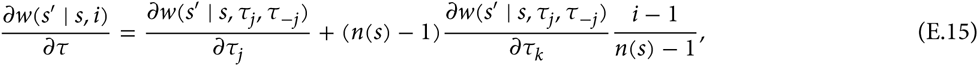

where *τ*_–*j*_ = (*τ*_1_, …, *τ*_*j*–1_, *τ*_*j*+1_, … *τ*_*n(s)* –1_) is the vector collecting the phenotypes of the neighbors of an individual *j* and *k ≠ j*. Substituting into eq. (E.14), setting 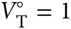, and using the definition of relatedness given in the Table 1 gives

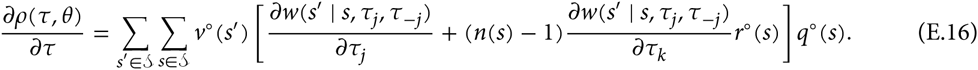

#### E.3.1 Connection to direct fitness method results

Here, we prove that eq. (E.16) returns exactly eqs. 26-27 of Rousset and Ronce (2004) when states are population sizes and each individuals migrates independently from each other. This proves that we recover in general the results obtained by the direct fitness method since the results of Rousset and Ronce (2004) generalize those of Taylor and Frank (1996) to demographically structured populations.

In order to show the connection, we need to prove that

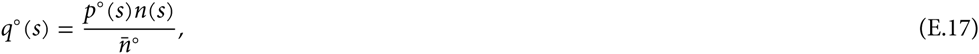

where 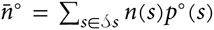 is the average group size in a monomorphic *θ* population. For this, we first note that from the definition of *q(s)* (Table 1), we have

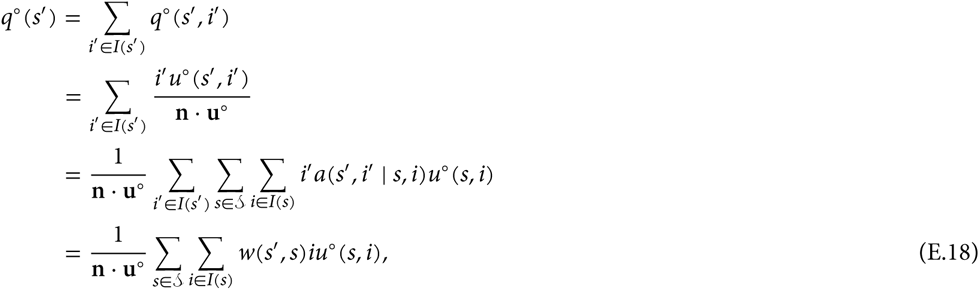

which yields

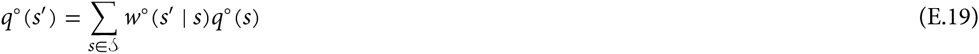

and shows that the vector collecting the *q*° (*s*) is a right unit eigenvector of the matrix with elements *w*° (*s*′ | *s*). Let us now substitute the trial solution 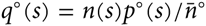 into eq. (E.19), whereby

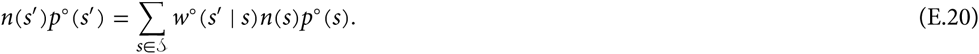

The right hand side is the total expected number of successful offspring in groups in state *s′* that descend from a randomly sampled group in the population. At stationarity this must be equal to *n(s′)p° (s′*), since *p° (s′)* is the probability of sampling a group in state *s′* and *n(s′)* is the number of successful offspring in that group. Hence, 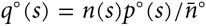 satisfies eq. (E.19) and eq. (E.17) holds.

We now expand eq. (E.16) by using the decomposition of personal fitness *w*(*s′* | *s, i*) = *w*_p_(*s′* | *s, i*) + *w*_p_ (*s′* | *s, i*) (eq. (6) of the main text), which allows us to write

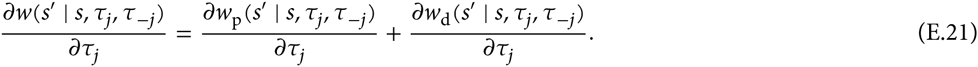

Each of these component will be further expanded by using eqs. (E.1)-(E.2). For the philopatric component, from eq. (E.1) we can write

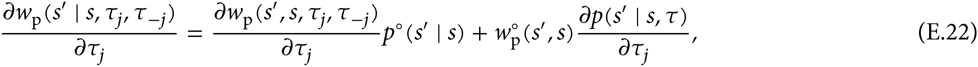

where *τ* = (*τ*_1_, …, *τ*_*n*,(*s*)–1_). For a neighbour *k ≠ j* of a focal mutant *j* we have

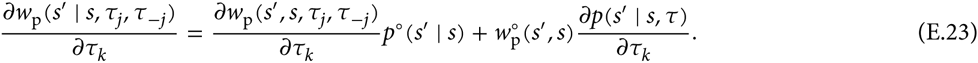

In order to expand the dispersal component in eq. (E.21), we follow the assumption of Rousset and Ronce, 2004 that the composition of a natal group of mutants does not affect the transition probability of other groups (owing to the fact that individuals migrate independently from each other) and set *k(s′* | *ζ, s, i*) = *p*° (*s′ | ζ*) in eq. (E.2). Then, we can write

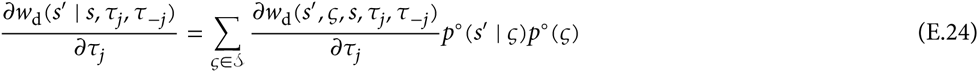

and for *k* ≠ *j*

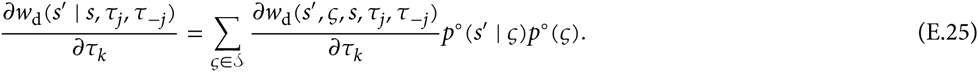

Substituting eqs. (E.21)–(E.25) into eq. (E.16) yields

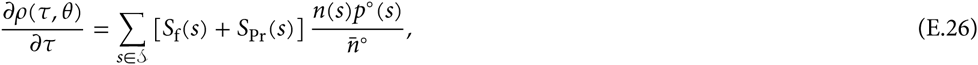

where

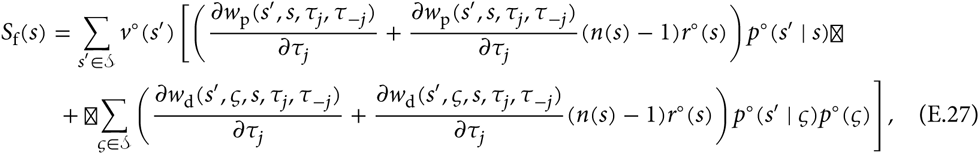

and

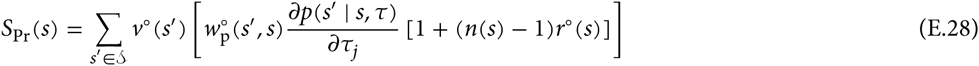

If we let *s* be group size and set *n(s)* = *s*, then eqs. (E.27)-(E.28) are proportional to eqs. (A.33)-(A.36) of Lehmann and Rousset (2010). If we multiply eq. (E.26) by *n(s′)/n(s′)* and use class reproductive values *α° (s′)* = *v° (s′)n(s′)* and the definition of frequency functions of Rousset and Ronce (2004, eqs. 33-34), then eqs. (E.27)-(E.28) are proportional to eqs. (26)-(27) of Rousset and Ronce (2004).

## Appendix F Fixed number of age or stage classes

We here consider a situation where there is a uniform demography, where each group is of constant size but now each individual belongs to one of a set of fixed classes where the set of class is given by *C* = {1, …, *n_c_*}. An example would age structure due to overlapping generations or different castes of social insects like workers and queens.

Let **i** = (*i*_1_, …, *i_n_c__)* ∊ *i* be the vector of the number of mutant alleles of type *τ* in class 1 to *n_c_* in a group where *i* is the set of possible configurations. Let *I* = (*I*_1_ × ⋯ × *I_n_c__*) **0** where *I_x_* = {0,1, …, *n_x_*} is set of the number of mutant alleles in class *x* and *n_x_* is the number of individuals in that class. We remove the all zero state **0** from *I* so that we only track states with at least one mutant in some class. Let **A** be the matrix with elements *a*(**i**′ | **i**) giving the expected number of groups in state **i**′ produced by a focal group in state **i**′. Further, let **n** be the vector collecting the total number of mutant individuals for each state; i.e., the i-th state of n is given by *x*(**i**) = Σ_*y∊C*_ *i_y_*.

We now prove the expression for lineage fitness (e.g., eq. 5) for this model and proceed in the same way as in Appendices A-C. Hence, we first note that

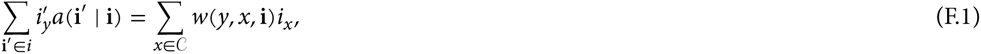

where *w*(*y, x*, **i**) is the expected number of class *y* offspring produced by a class *x* mutant when in a group in state **i**. Now, from **n** · *ρ***u** = (**n** · **Au**)and eq. (F.1), we have

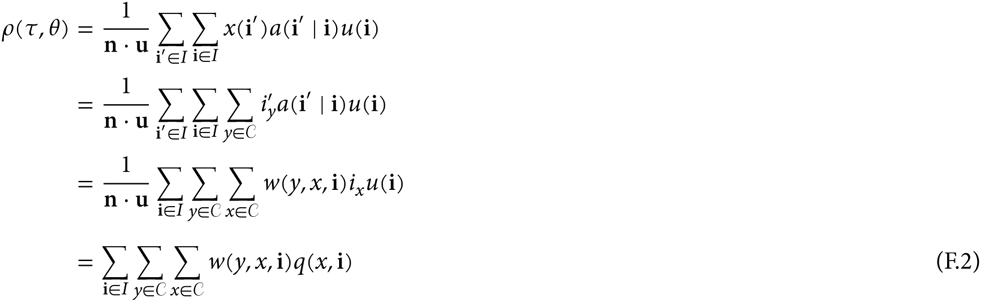

where

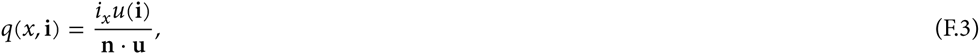

which satisfies Σ_*x∊C*_ Σ_x∊*I*_ *q_x_*(**i**) = 1. Defining lineage fitness as

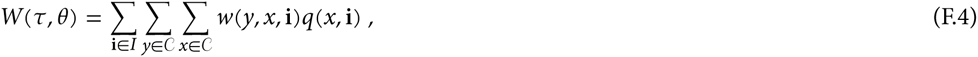

eq. (F.2) shows that *ρ(τ, θ)* = *W*(*τ, θ*), which is the same result as eq. (7).

Second, we derive an expression for the inclusive fitness *W*_IF_ (*τ, θ*). Inclusive fitness requires that we calculate reproductive values, so we gather into the vector **v**° the elements

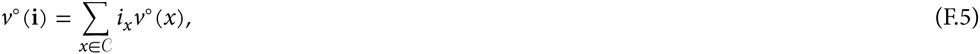

where *v*° (*x*) is the reproductive value of an individual in class *x*. Now, from **v**° · *ρ***u** = (**v**° · **Au**) and eq. (F.5),
we have

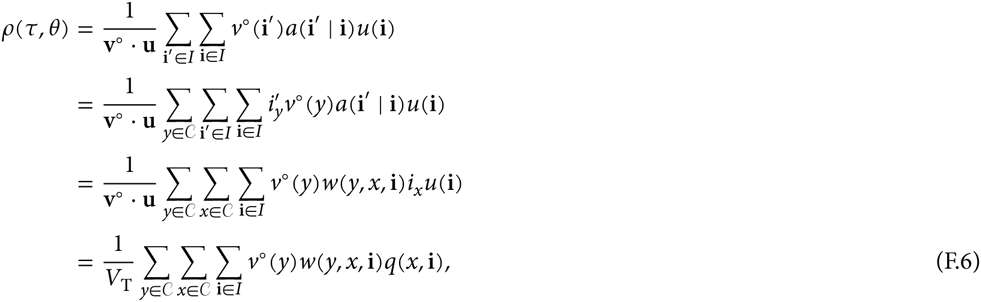

where

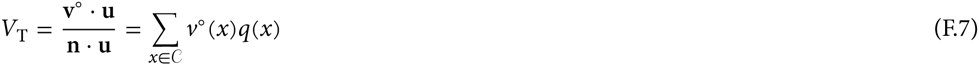

and *q(x)* = Σ_i*∊I*_ *q*(*x*, **i**) is the probability of sampling a lineage member in class *x*. Suppose we now form a weighted multiple regression

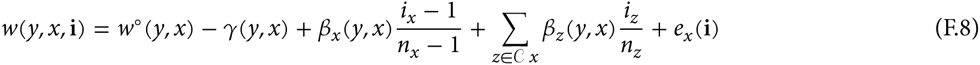

and least square fit *γ* and the *β*’s by minimizing

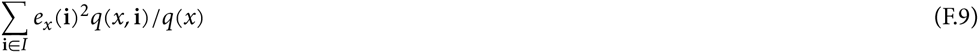

where the weights are given by *q_x_*(**i**)/*q_x_*. This procedure guarantees that the weighted sum of errors is zero, or that *Σ_i*∊I*_ = *e_x_* (**i**)(*q_x_*(**i**)/*q_x_*) =* 0. Let us further define

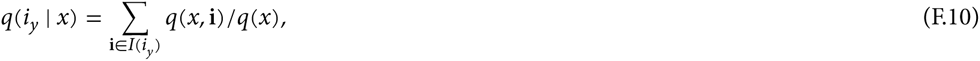

where *I*(*i_y_*) denotes the elements of the set *I* whose number of class *y* mutants is equal to *i_y_*. Then, we can interpret *q*(*i_y_* | *x*) as the probability that there are *i_y_* mutants in class *y* given that a mutant has a sampled a mutant in class *x*. Substituting all this into eq. (F.6), we have

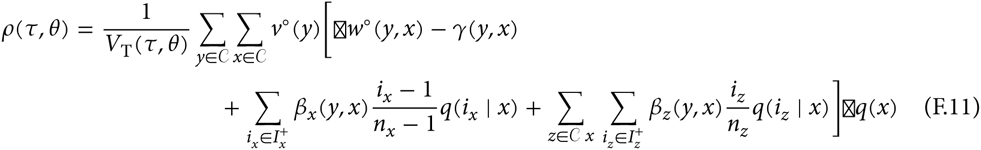

where we only sum over the elements 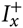 = {1, …, *n_x_*} in the first sum of the second line since *w*(*y, x*, **i**) = 0 for all *i_x_* = 0 (the second sum in the second line uses 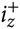 for ease of notation).

Let us now define inclusive fitness as

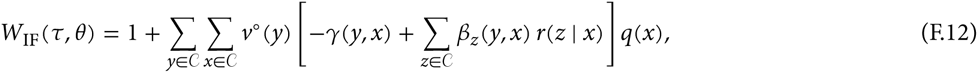

where

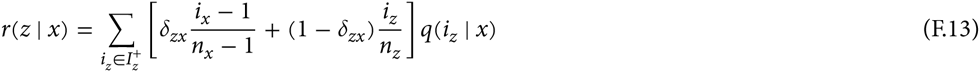

is the probability that, conditional on being sampled in class *x*, an individual carrying the mutant experiences a randomly sampled neighbour in class *z* that also carries the mutant allele, and where *δ_zx_* is the Kronecker Delta (*δ_zx_* = 1 if *z* = *x*, zero otherwise). Substituting eqs. (F.12)-(F.13) into eq. (F.11) and using the definition of reproductive value we obtain

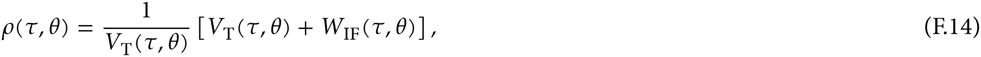

whereby

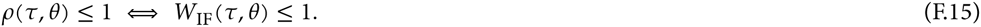

